# A high-level programming language for generative protein design

**DOI:** 10.1101/2022.12.21.521526

**Authors:** Brian Hie, Salvatore Candido, Zeming Lin, Ori Kabeli, Roshan Rao, Nikita Smetanin, Tom Sercu, Alexander Rives

## Abstract

Combining a basic set of building blocks into more complex forms is a universal design principle. Most protein designs have proceeded from a manual bottom-up approach using parts created by nature, but top-down design of proteins is fundamentally hard due to biological complexity. We demonstrate how the modularity and programmability long sought for protein design can be realized through generative artificial intelligence. Advanced protein language models demonstrate emergent learning of atomic resolution structure and protein design principles. We leverage these developments to enable the programmable design of de novo protein sequences and structures of high complexity. First, we describe a high-level programming language based on modular building blocks that allows a designer to easily compose a set of desired properties. We then develop an energy-based generative model, built on atomic resolution structure prediction with a language model, that realizes all-atom structure designs that have the programmed properties. Designing a diverse set of specifications, including constraints on atomic coordinates, secondary structure, symmetry, and multimerization, demonstrates the generality and controllability of the approach. Enumerating constraints at increasing levels of hierarchical complexity shows that the approach can access a combinatorially large design space.

## Introduction

Protein design would benefit from the regularity, simplicity, and programmability provided by a basic set of abstractions (1–4) like those used in the engineering of buildings, machines, circuits, and computer software. But unlike these artificial creations, proteins cannot be decomposed into easily recombinable parts because the local structure of the sequence is entangled in its global context (5, 6). Classical de novo protein design has attempted to determine a fundamental set of structural building blocks, which could then be assembled into higher-order structures (7–11). Likewise, traditional protein engineering often recombines segments or domains of natural protein sequences into hybrid chimeras (12–14). However, existing approaches have not been able to achieve the high combinatorial complexity that is necessary for true programmability.

We show modern generative models realize these classical goals of modularity and programmability at a new level of combinatorial complexity. Our idea is to place the modularity and programmability at a higher level of abstraction, where a generative model bridges the gap between human intuition and the production of specific sequences and structures. In this setting, the protein designer needs only to recombine high-level directives, while the task of obtaining a protein that fulfills those directives is placed on the generative model.

We propose a programming language for generative protein design, which allows a designer to specify intuitive, modular, and hierarchical programs. We show that high-level programs can be translated into low-level sequences and structures by a generative model. Our approach leverages advances in protein language models, which learn structural information (15, 16) and the design principles of proteins (see accompanying paper by Verkuil et al.).

In this study, our specific implementation is based on an energy-based generative model. First, a protein designer specifies a high-level program consisting of a set of hierarchically organized constraints (Figure 1A). Then, this program compiles to an energy function that evaluates compatibility with the constraints, which can be arbitrary and non-differentiable (Figure 1B). We apply constraints on structure by incorporating atomic-level structure predictions, enabled by a language model, into the energy function. This approach enables the generation of a wide set of complex designs (Figure 1C).

**Figure 1.**
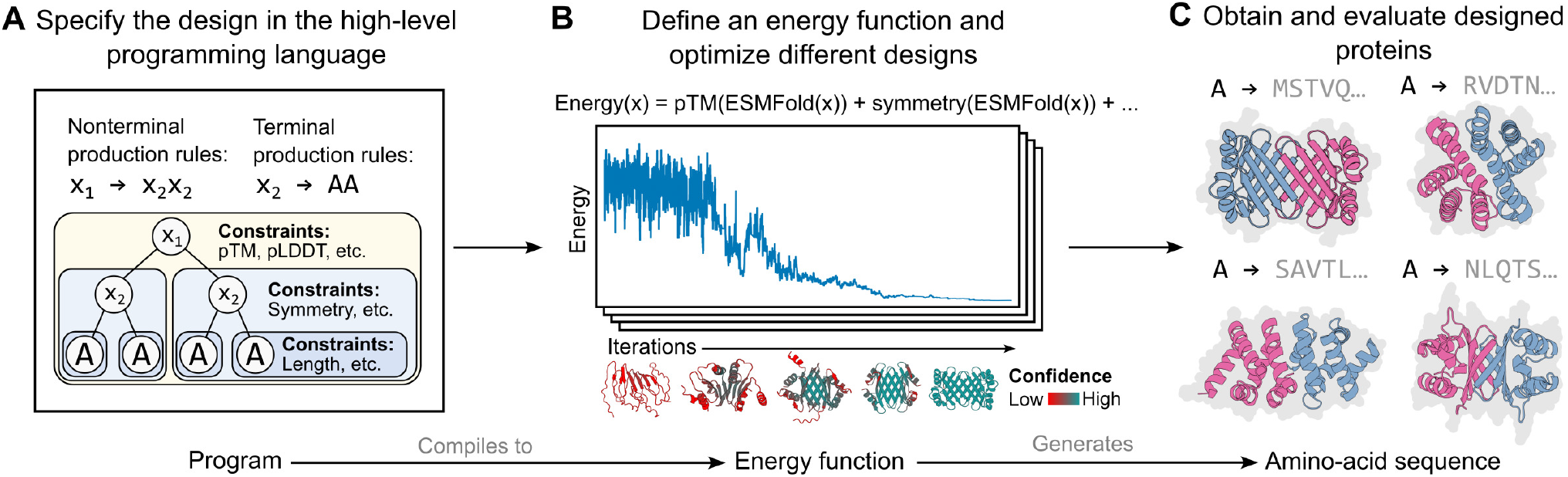
Overview of the high-level programming language and the optimization algorithm. **(A)** We propose a high-level programming language in which each program consists of (1) a syntax tree (corresponding to a set of nonterminal and terminal production rules) that enables modular and hierarchical organization of protein subunits and (2) a set of constraint functions that can be defined at each node of the syntax tree, where a given constraint is applied to the entire subtree rooted at the corresponding node. **(B)** This program is then compiled to a single energy function, which in our study is a simple linear combination of the specified constraint functions. The energy function is used to guide an optimization procedure based on simulated annealing, of which a key component is the use of an accurate and efficient structure predictor to evaluate the energy function at each step of the optimization. The same energy function can guide multiple optimization trajectories. **(C)** Each of these trajectories produces a protein sequence design and an associated predicted structure. These sequences and predicted structures can then be evaluated downstream using in silico and experimental metrics.

The use of a high-level language allows the protein designer to systematically reason about the design space and specify very general, modular, and composable programs. To demonstrate this, we generate proteins that realize a variety of constraints that include secondary structure, symmetry, multimerization, and atomic-level coordination in the predicted structures.We apply these constraints in complex, hierarchical settings, where we can enumerate a space of highly idealized forms that have low similarity to natural structures. As de novo design progresses to more complex proteins and protein assemblies, high-level abstractions such as the programming language described in this study should facilitate the systematic exploration and design of complex artificial proteins.

### A generative programming language for protein design

We introduce a high-level programming language for generative protein design. This language first requires a syntax tree (Figure 1A) consisting of terminal symbols (i.e., the leaves of the tree) that each corresponds to a unique protein sequence (which is potentially repeated within the protein) and nonterminal symbols (i.e., the internal nodes of the tree) that enable hierarchical organization. Second, the language requires a set of constraints: at each node in the tree, a protein designer can specify any number of constraints, which are applied to the entire subtree. The syntax tree and its constraints fully specify a program in our high-level language. We provide a more extended description of this language in the Methods section.

Each program is compiled into an energy function that specifies a generative model for that program in the form of a distribution

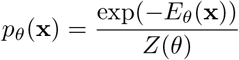

over protein sequences x conforming to the program. The constraints are encoded as weighted terms that are additively combined into the total energy. Since the partition function *Z*, which is a function of the parameters *θ*, is intractable, low temperature samples can be taken with MCMC and simulated annealing (Figure 1B). The generative capacity of this approach is built on recent developments in deep learning for protein biology. Specifically, each step in the optimization loop has access to a fast and accurate atomic-level structure prediction enabled by the ESM-2 protein language model.

Given a single program, the generative model can create potentially diverse designs that fulfill the user-specified constraints (Figure 1C). These constraints can be arbitary and nondifferentiable, and can span multiple scales of biological complexity, from atomic-level coordinates to abstract plans of the protein including the overall topology and symmetry. This approach allows the model to propose diverse solutions where many potential designs may satisfy the program. By leveraging an expressive model of structure in a generative capacity, the resulting designs respect the various constraints individually and are also globally coherent.

## Results

### Full-protein constraints

We first demonstrate that our approach can design proteins where the constraints are simply applied to the entire sequence and structure without any hierarchical organization. An especially valuable constraint, which we apply generally across all our design efforts, steers the optimization toward predicted structures with higher model confidence, i.e., high pTM and mean pLDDT. For proteins that we desire to have soluble, monomeric expression, we also steer the optimization to minimize hydrophobic residues that are solvent-exposed (Methods). Using only these constraints on structural confidence and hydrophobic residue placement, our model is able to generate or “freely hallucinate” (17) high-confidence structures (Figures 2A and 2B); across 200 seeds, all optimization loops produced predicted structures with an ESMFold mean pLDDT greater than 0.7 (Figure 2B). Of these, a large portion (44, or 22%) also had high predicted confidence (pLDDT *>* 0.7) by single-sequence AlphaFold2 (18) (Figure 2C), a separate structure prediction model that was not used in our optimization procedure.

**Figure 2.**
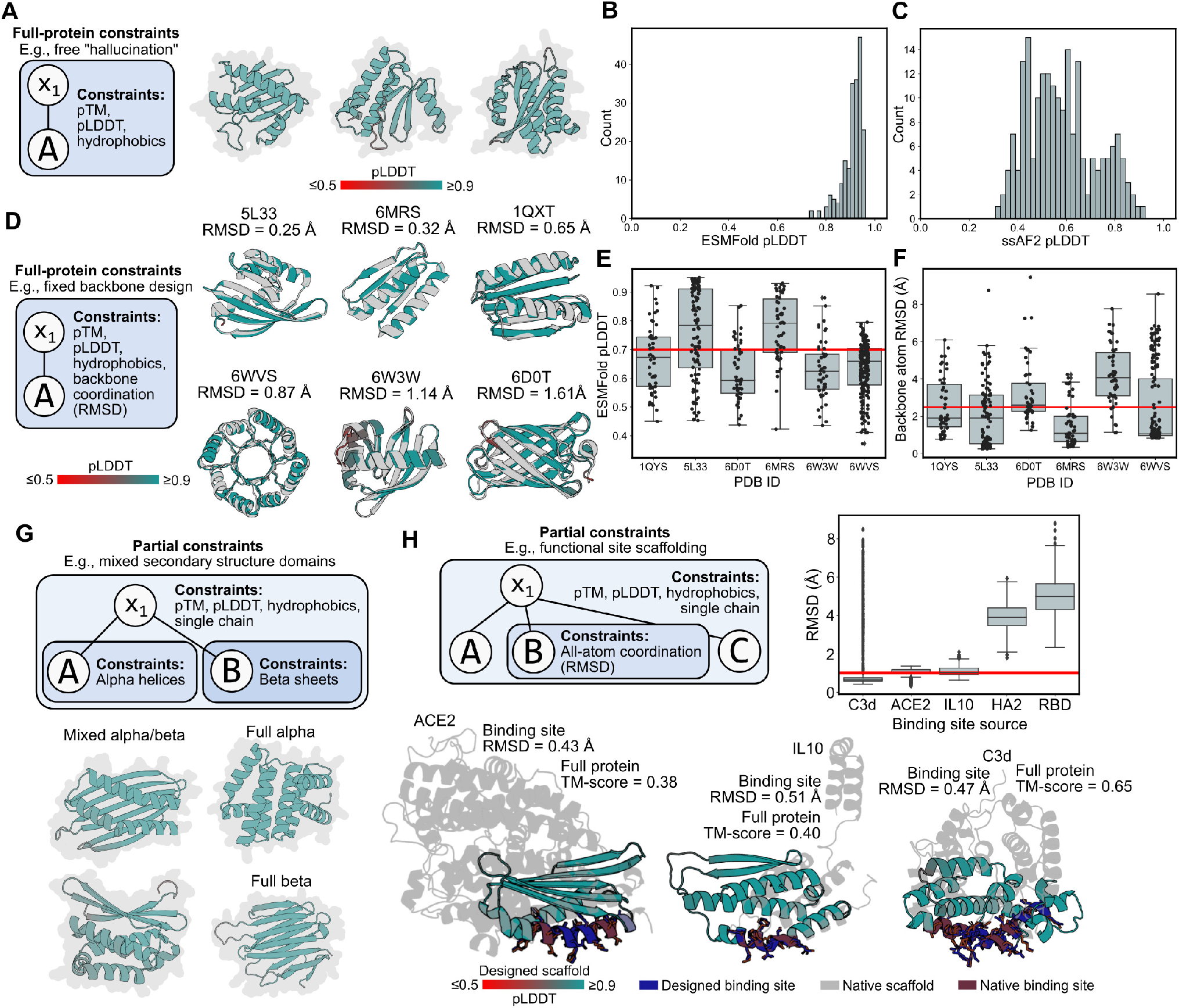
Programming full-protein or partial constraints. **(A)** A graphical representation of a program for protein “free hallucination” (left) along with three example designed structures (right). **(B)** The distribution of ESMFold pLDDT values over 200 free-hallucinated structures. Of these, 100% have good confidence (ESMFold pLDDT > 0.7). **(C)** The distribution of single-sequence AlphaFold2 (ssAF2) over the same 200 structures; note that ssAF2 was not used in the design procedure. Of these, 22% have good confidence (ssAF2 pLDDT > 0.7). **(D)** A graphical representation of a program for fixed backbone design (left) along with example designs for six de novo target backbones. The experimental backbone is colored gray; the designed backbone is colored by ESMFold pLDDT. **(E)** For each target backbone, the distribution of the ESMFold pLDDT values of the final designs from 50 or more fixed backbone design seeds is plotted as a boxplot (for all boxplots in this figure, the box extends from first to third quartile, black line indicates the median, and whiskers indicate 2.5 times the interquartile range) with each seed also plotted as a black circle. A horizontal red line indicates pLDDT = 0.7. For each target backbone, the distribution of RMSD values between the target and design backbone atoms from 50 or more fixed backbone design seeds is plotted as a boxplot with each seed also plotted as a black circle. A horizontal red line indicates RMSD = 2.5 Å. A graphical representation of a program for designing a protein with mixed secondary structure (top) along with example designs in which secondary structure was explicitly specified (bottom). **(H)** Top-left: A graphical representation of a program for functional site scaffolding. Top-right: For each scaffolded binding site, the distribution of RMSD between the native and designed binding site atoms (including side chains) from 2,000 seeds is plotted as a boxplot. A horizontal red line indicates RMSD = 2 Å. Bottom: Example designs that achieve sub-angstrom atomic coordination in the scaffolded binding site atoms, high model confidence in the associated scaffold, and low similarity (quantified by TM-score) to the natural protein.

Our objective function also enables other full-protein constraints, such as specifying the positions of the backbone atoms while allowing the algorithm to design the corresponding sequence, a design task referred to as fixed backbone design (19). To achieve this, we can add a term to the energy function that minimizes the root-mean-square deviation (RMSD) between the corresponding designed and target backbone atoms (Methods). Our simulated annealing procedure successfully produces high-confidence designs with low RMSD (< 1.6 Å) across diverse de novo backbones (Figures 2D and 2E), and can do so reproducibly over different optimization runs (Figure 2F).

### Partial constraints

We next sought to increase the complexity of our designable space by varying the constraints enforced on different parts of a protein. For example, a simple mixed-constraint setting is to specify a two-domain protein with different combinations of secondary structure composition (Figures 2G and S1A–S1C). In our programs, we can represent this setting by a syntax tree containing two or more subtrees, where different constraints are only applied within the discrete subtrees.

A more complex mixed-constraint setting is to design functional proteins by constraining one region of the protein design to have the same all-atom positions (including protein side chains) as a functional site from nature, while allowing the design procedure to freely generate the remainder of the protein; this design setting is sometimes referred to as functional site “scaffolding” (20). Importantly, in contrast to fixed backbone design, in which constraints are only placed on backbone atomic coordinates, functional site scaffolding requires constraints on side-chain atoms as well, since these are critical to achieving function. Because our optimization procedure produces an all-atom structure prediction at each step of the optimization, we can readily incorporate this constraint as part of the energy function by minimizing the all-atom RMSD between the natural and designed atomic coordinates of the functional site (Methods).

Across functional sites involving sequence-contiguous or -discontiguous residues from a variety of natural proteins, our algorithm is able to produce designs that scaffold the site with sub-angstrom RMSD between the experimental and predicted structure in three out of five functional sites attempted (Figures 2H and S1D). Moreover, the algorithm produces designed scaffolds that depart from the native protein (Figure 2H). The ability to move natural functional sites onto designed backbones has many practical applications, including the design of functional proteins that are smaller or stabler than their natural counterparts.

### Symmetric and multimeric group constraints

Beyond proteins containing partial constraints, we next increase the complexity of our protein designs by generating structures that contain constraints over multiple subunits. A foundational design task for the generation of idealized, de novo proteins is to constrain structural symmetry (7, 22). To generate symmetric proteins, we first enforce the notion of a repeated unit that is repeated *K* times when designing a *K*-fold symmetry (where we can control the value of *K*). To guide the optimization toward symmetric structures, we add various constraints on the distances among the centroids of each repeated unit as part of the energy function (Methods). In our high-level language, a symmetric protein would be encoded by repeating the same non-terminal symbol *K* times (corresponding to the repeated unit); the symmetry constraint is then placed at the level of the syntax tree containing these repeated non-terminals (Figure 3A).

**Figure 3.**
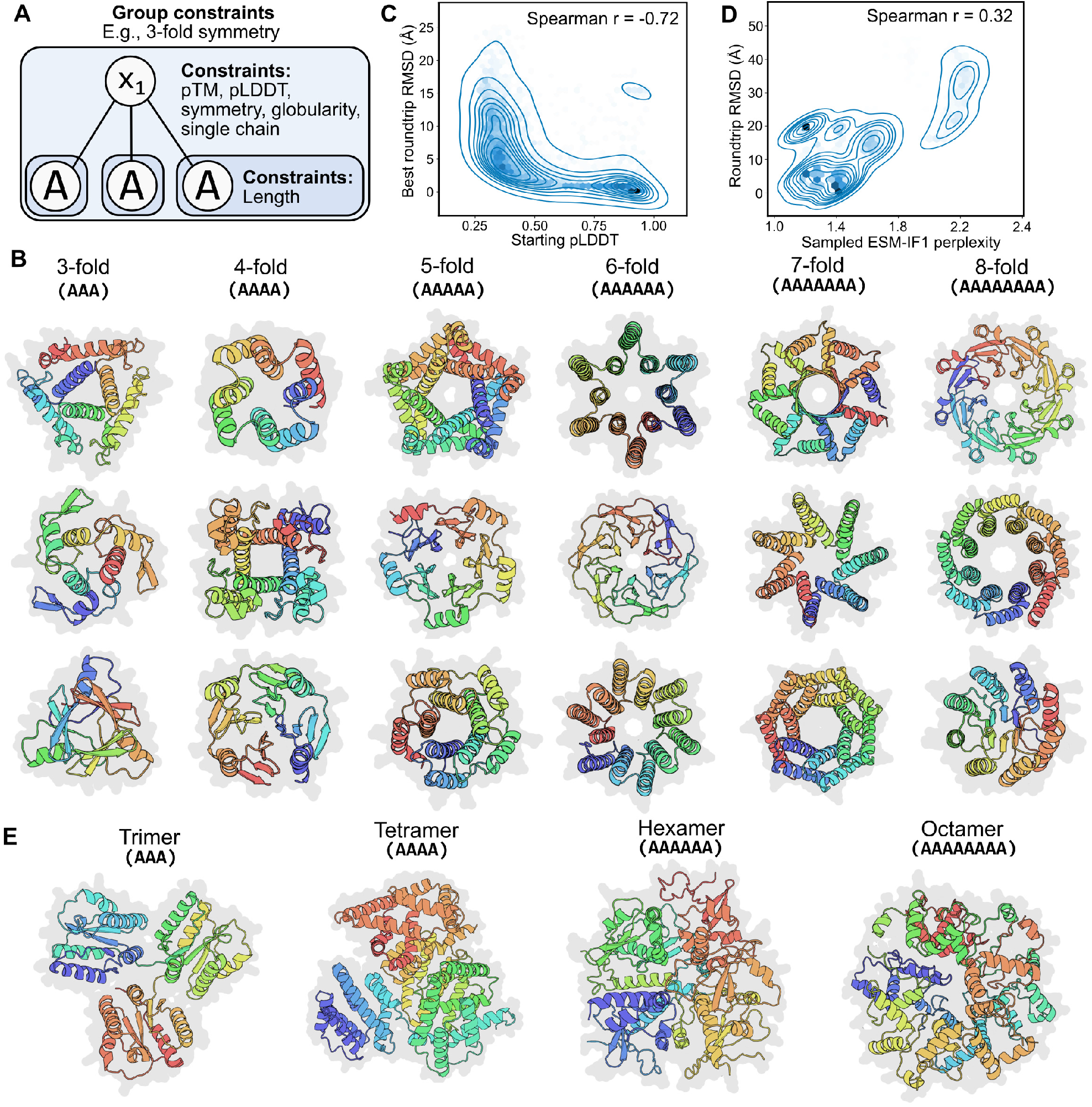
Programming symmetry and homo-oligomerization. **(A)** A graphical representation of a program for designing a single protein chain with 3-fold symmetry based on a repeated subsequence. **(B)** Example designs varying fold symmetry from 3-to 8-fold. **(C)** 1000 randomly sampled symmetric protein designs were “roundtripped” by sampling ten sequences via ESM-IF1 inverse folding (21) of their backbones followed by ESMFold structure prediction. The ESMFold pLDDT of the starting backbone is indicated on the horizontal axis. The lowest of the 10 RMSDs comparing the starting and roundtripped backbone atoms is indicated on the vertical axis. Blue lines indicate density contours and hexagonal bins are darker with greater density. We observed that a more confident design is associated with roundtrip success. **(D)** 1,000 randomly sampled inverse folding samples are plotted according to their ESM-IF1 perplexity on the horizontal axis and their roundtrip RMSD on the vertical axis. We observed that a lower perplexity sequence is associated with roundtrip success. **(E)** Example homo-oligomers with increasing numbers of individual protomers. The tetrameric, hexameric, and octameric oligomers depicted here form globular polyhedral shapes rather than the rotational symmetry of designs in **(B)**.

Using these symmetric constraints, we show that we can program the level of symmetry within a protein design. When directed to design 3-to 8-fold symmetry, the generative model produces a diverse set of high-confidence structures (Figures 3B, S2A, and S2B), including folds that have common analogs in nature (including coiled-coils, beta propellers, beta sandwiches, beta barrels, and TIM barrels) as well as highly-idealized designs that are different from natural structures, including a pentagonal star-shaped protein (with a TM-score of 0.48 to the nearest PDB structure 3S38; row 1 and column 3 in Figure 3B) and a cube-shape protein (nearest-PDB TM-score of 0.51 to PDB 7DEG; row 2 and column 2 in Figure 3B). The highlighted symmetric proteins in Figure 3B have nearest-PDB TM-scores ranging from 0.47 to 0.86, with a median TM-score of 0.64; TM-scores for all seeds are plotted in Figure S2C.

To increase our confidence that these idealized structures correspond to valid and designable backbones, we observe that sampling sequences via inverse folding with the ESM-IF1 and ProteinMPNN models (21, 23) followed by structure prediction of the sequence samples can reproducibly recover the original backbone geometry (Methods), in many instances with sub-angstrom backbone-atom RMSD. To increase our confidence that a successful “roundtrip” through inverse folding indicates designable backbones, we observe that high confidence designs indicates better roundtrip success (Figures 3C and S2D) and that the ability to sample a low perplexity sequence also indicates roundtrip success (Figure 3D).

Beyond single-chain symmetries, we can also design multimeric proteins similarly. We enforce the notion of single or multiple chains in our programming language with a “single chain” constraint that dictates that all terminal elements in a subtree belong to the same chain (Methods); to design multimeric proteins, we need only remove this constraint. Example multimeric symmetric proteins involving 4-to 8-mers are provided in Figure 3E.

### Hierarchical constraints

Formalizing our constraints into a syntax tree naturally enables the specification of hierarchical constraints, which enables more complex protein designs. As an initial demonstration, guided by our high-level language’s formalization, we design different levels of symmetry at two levels of hierarchy, where the lower level of symmetry is specified within a chain and the upper level of symmetry is specified among protomers in a homo-oligomer (Figure 4A). We procedurally enumerate over examples that range from a dimer of units with 2-fold symmetry to a tetramer of units with 4-fold symmetry (Figures 4B, S3A, and S3B). As in the single-chain symmetric design setting, we observe successful structure prediction roundtrips through inverse folding, and that both high-confidence predicted structures and low inverse folding perplexity indicate roundtrip success (Figures 4C and 4D). Many of the designs with two levels of symmetry have low overall similarity to structures in the PDB; for example, a dimer of 2-fold symmetry in which opposing beta sheets form a regular checkerboard pattern (nearest-PDB TM-score of 0.49 to PDB 3W38; row 1 and column 1 in Figure 4B). The highlighted homo-oligomers of two-level symmetry in Figure 4B have nearest-PDB TM-scores ranging from 0.25 to 0.52, with a median TM-score of 0.48; TM-scores for all seeds are plotted in Figure S3C.

**Figure 4.**
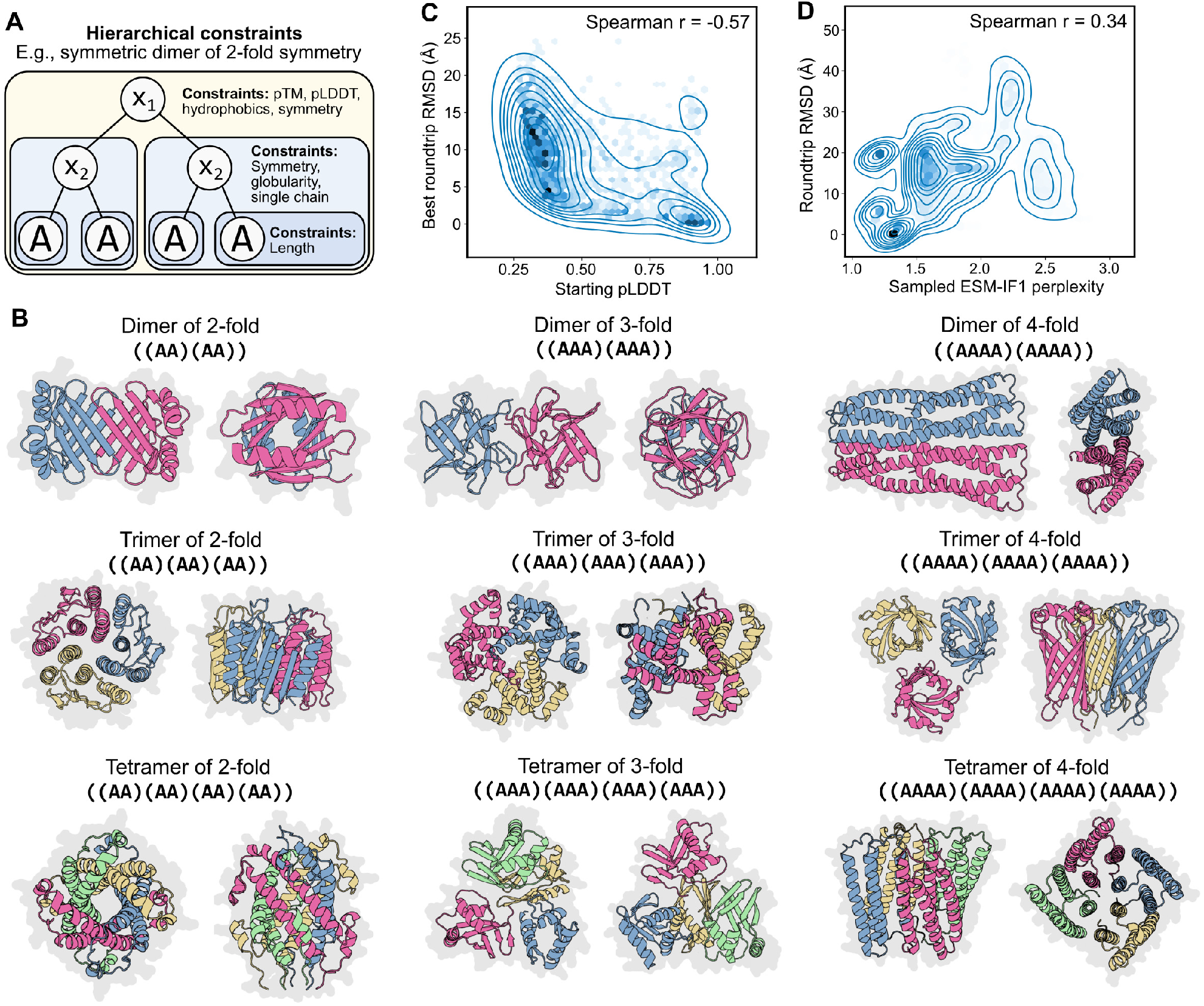
Programming two levels of symmetry. **(A)** A graphical representation of a program for designing two levels of symmetry in which a homo-oligomeric symmetric dimer represents the top level of symmetry and each unit within the dimer also has two-fold symmetry. **(B)** Example oligomers with two levels of symmetry, in which we procedurally enumerate across a grid in which we vary the top-level symmetry across the rows and the bottom-level symmetry across the columns. Discrete chains are indicated by different colors. **(C)** 1,000 randomly sampled two-level symmetric protein oligomer designs were “roundtripped” by sampling ten sequences via ESM-IF1 inverse folding (21) of their backbones followed by ESMFold structure prediction (Methods). The ESMFold pLDDT of the starting backbone is indicated on the horizontal axis. The lowest of the 10 RMSDs comparing the starting and roundtripped backbone atoms is indicated on the vertical axis. Blue lines indicate density contours and hexagonal bins are darker with greater density. We observed that a more confident design is associated with roundtrip success. **(D)** 1,000 randomly sampled inverse folding samples are plotted according to their ESM-IF1 perplexity on the horizontal axis and their roundtrip RMSD on the vertical axis. We observed that a lower perplexity sequence is associated with roundtrip success.

Another hierarchical design setting is to combine the function-scaffolding and the symmetric design tasks described above, as some functions are enhanced by repetition of a functional site; for example, when improving the strength of a binding interaction, multiple binding sites on a protein could synergize such that the overall binding avidity is greater than the sum of the individual affinities (24). This task requires two levels of hierarchy: the top level specifies symmetry while the bottom level specifies the side-chain atomic coordination constraint (Figure 5A). With this corresponding program, we can generate designs in which an atomic-level constraint is enforced on multiple functional sites over the protein, the overall protein organization is constrained to be symmetric, and we can control the level of designed symmetry (Figures 5B and S4A–S4C).

**Figure 5.**
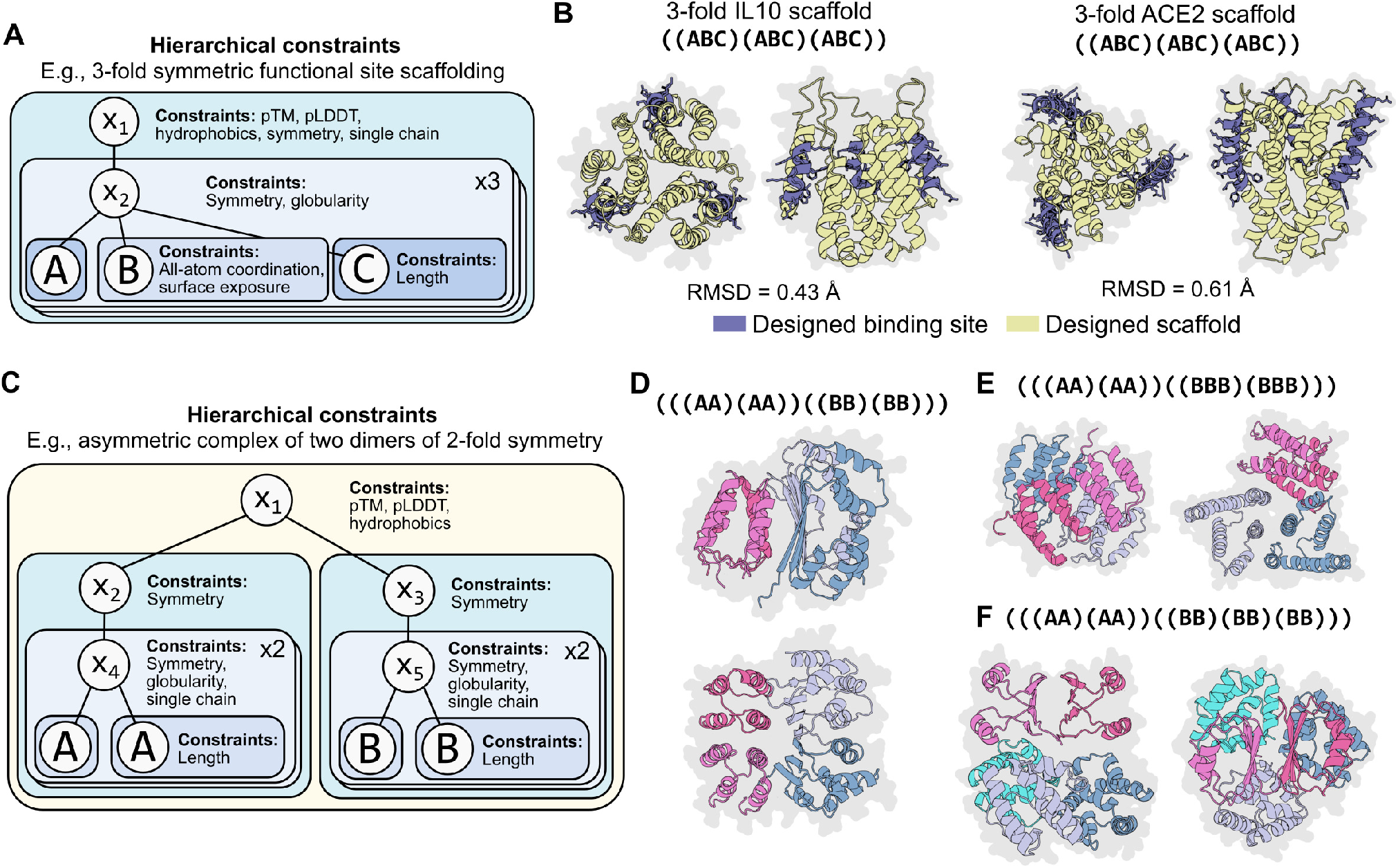
Programming complex hierarchical constraints. **(A)** A graphical representation of a program for scaffolding three functional sites in which those sites have a 3-fold symmetry. **(B)** Example 3-fold symmetric scaffolds for the IL10 and ACE2 binding sites that achieve sub-angstrom RMSD averaged across the three sites. **(C)** A graphical representation of a program that specifies an asymmetric protein complex consisting of two pairs of chains. Each pair is constrained to have 2-fold symmetry between the constituent chains. Furthermore, each constituent chain itself has 2-fold symmetry. **(D)** A generated protein structure as specified by the program depicted in **(C)**. Discrete chains are indicated by different colors. **(E)** A generated protein structure as specified by the program depicted in **(C)** except where one of the pairs has constituent chains that have three-fold symmetry. Discrete chains are indicated by different colors. **(F)** A generated protein structure as specified by the program depicted in **(C)** except where one of the pairs is replaced with a symmetric trimer (where each constituent chain in the trimer has two-fold symmetry). Discrete chains are indicated by different colors.

We lastly show that we can specify protein designs that have even deeper levels of hierarchy in their constraints (Figure 5C) by designing protein assemblies that combine both symmetry and asymmetry. For example, we designed a protein complex composed of four units in which a pair of the chains are symmetric to each other (and each unit internally has two-fold symmetry) and where another pair of chains are symmetric to each other (and each unit also internally has two-fold symmetry), but the two pairs are asymmetric to each other (Figures 5C, 5D, S4D, and S4E). Our high-level programming language readily enables us to control the complexity of the generated complexes such that, for example, one of the pairs consists of chains with threefold symmetry (Figure 5E) or that the complex consists of five chains (a pair of symmetric chains of two-fold symmetry asymmetrically complexed with a triple of symmetric chains of two-fold symmetry) (Figure 5F). We find that our optimization procedure can produce designed structures consistent with all of these hierarchical specifications.

## Related work

This paper is related to classical work that attempts to (i) classify a set of common sequence or structure motifs (3, 4) and (ii) manually combine these motifs to generate new proteins (7–14). More recently, deep-learning-based methods have increased the complexity of designable structures (17, 20, 22) and machine-learning-based generative models have shown increasingly sophisticated design capabilities. These include sequence-based Potts models and autoregressive language models for designing sequences (25–27), Markov Chain Monte Carlo algorithms combined with structure prediction for jointly designing sequences and structures (17, 20, 22), inverse folding models that use structural backbone coordinates to design sequences (21, 23), and concurrent work using diffusion models for designing protein backbones (28, 29). A key contribution of this study is to combine the modularity aspired to by classical methods with the power of modern generative models, in particular improvements in the accuracy and efficiency of language-model-based protein structure prediction (16).

## Discussion

In this study, we show that generative artificial intelligence enables high-level programmability at a new level of combinatorial complexity. We propose a programming language that can express high-level programs for the design of proteins at diverse biological scales, including atomic-level coordinates, secondary structure, and high-level symmetries within single chains and the units of self-assembling multichain complexes. We show that programs written in the abstract language can be compiled into an energy function and that the corresponding generative model is capable of fulfilling complex constraints within an overall coherent structure.

We demonstrate programs of increasing levels of complexity, including the design of homo-oligomers with two levels of symmetry, symmetric functional scaffolds, and asymmetric complexes of subunits that themselves have two levels of symmetry. The approach reveals a large space of idealized protein designs created from top-down design principles. Especially as the complexity of the constraints increases, many of the corresponding designs are highly idealized, analogous to the regularity of artificially created machines and systems.

Our computational results using two independent inverse folding methods suggest that the generated structures are designable, since inverse folding models have demonstrated high experimental success rates (23). We are also obtaining data to experimentally validate the designs.

More broadly, the formalization offered by a high-level programming language enables logical design principles to be applied to protein design as in other fields of engineering. This has been especially challenging in biology due to the way that the amino acid sequence opaquely encodes the structure and function. As protein design moves toward the engineering of more complex functions, we anticipate that such a system will become increasingly useful.

## Acknowledgements

We would like to thank Halil Akin, Adam Lerer, Wenting Lu, Yaniv Shmueli, Robert Verkuil, and Zhongkai Zhu for technical help, feedback, and discussions that helped shape this project. We thank Laurens van der Maaten, Ammar Rizvi, Jon Shepard, and Joe Spisak for program support.

## A. Methods

### A.1 High-level programming language and energy-based optimization

A program in our language is fully specified by (1) a syntax tree and (2) a set of constraints. This program compiles to an energy function, which is used to guide black-box optimization of a protein sequence while also leveraging its predicted structure.

#### A.1.1. Syntax tree

The syntax tree consists of nonterminal symbols, which we denote as *x*_*i*_, as well as terminal symbols, which in our examples we denote as uppercase alphabetic characters such as *A, B, C*, etc. Each terminal symbol defines a unique protein sequence. The nonterminal symbol *x*_1_ is designated as the special start symbol; all programs must have *x*_1_. Additional nonterminal symbols are used to define hierarchical complexity. For example, specifying two levels of hierarchy requires a nonterminal production rule in addition to a terminal production rule, for example,

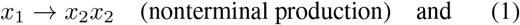

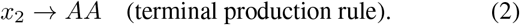

In the example above, the *x*_2_ nonterminal enables an intermediate level of hierarchy. A nonterminal can produce any finite-length permutation of higher-numbered nonterminals (for example, *x*_1_ → *x*_2_*x*_3_ is permitted but not *x*_2_ → *x*_1_*x*_3_ or *x*_2_ → *x*_2_*x*_3_). A nonterminal can also produce any finite-length permutation of terminals (for example, *x*_1_ → *AB*) or any finite-length permutation of mixed terminals and higher-numbered nonterminals (for example, *x*_1_ → *x*_2_*B* or *x*_1_ → *Bx*_2_).

A complete syntax tree is built by fully expanding the non-terminal *x*_1_ into a set of terminals. The production rules define the connectivity structure of the tree, where the parent node corresponds to the left side of the production rule and the child node(s) corresponds to the right side of the production rule. Across the entire syntax tree, each internal node corresponds to a nonterminal symbol and each leaf corresponds to a terminal symbol. Example syntax trees are provided in the main text figures.

#### A.1.2. Constraints

A program in our language also requires a set of constraints, where a single constraint is defined with respect to a single node and all of its descendants in the syntax tree. Note that this includes constraints on the leaves of the tree (corresponding to the terminal symbols). More specifically, a constraint is a function that takes as input the (sub)tree, as well as its corresponding (sub)sequence and (sub)structure, and outputs a real number. For example, a constraint defined with respect to the node corresponding to *x*_1_ simply receives as input the entire syntax tree, the full-length sequence, and the full protein structure. The same constraint (i.e., the same function) can be applied to multiple nodes in the tree. We will use *f*_*j*_(*x*_*i*_) to denote a constraint *j* defined with respect to the *x*_*i*_ node.

#### A.1.3. Compilation of constraints into an energy function

We compile a program into an energy function. In our study, we simply compute a linear combination of all the constraints in the user-specified set, i.e., *E*(*x*) = ∑_*i*_ ∑_*j*_ *f*_*j*_(*x*_*i*_), where *f*_*j*_(*x*_*i*_) is defined as zero when a constraint is not applied to a given node. In practice, we explicitly keep track of a scalar multiplicative weight on each constraint, i.e., *E*(*x*) = ∑_*i*_ ∑_*j*_ *a*_*j*_*f*_*j*_(*x*_*i*_). This energy is used in the simulated annealing optimization procedure described below. Specific examples of constraint functions used in our study are also provided below.

Linear combinations work well for our choice of generative model, but in principle any combination of the energy terms could be used here. For example, if we compiled our program into an energy function for a generative model that used a reward function (like a reinforcement learning agent), we might prefer a multiplicative combination of the inverse of our current energy terms.

#### A.1.4. Simulated annealing

The energy function is used as part of an iterative black-box optimization loop, where over multiple iterations, a change to a given state (in this case, a protein sequence design) is accepted with some probability. We use a simulated annealing algorithm in which the acceptance probability is controlled by a temperature value such that the optimization can tolerate higher energy changes at the beginning of the optimization before favoring changes that decrease the energy toward the end of the optimization. In our study, we begin by initializing the sequence state (one unique sequence per terminal symbol) with uniform amino-acid probability to a given user-specified length; we also compute an initial structure prediction from this sequence.

Each iteration proposes a mutation to the protein sequence. To make this proposal, first, one of the terminal symbols is chosen with uniform probability, and second, one of a substitution, insertion, or deletion is chosen with some probability (we default to 60%, 20%, and 20%, respectively). For substitutions and insertions, the new amino acid is chosen with some probability (unless otherwise specified, we apply uniform probability over a reduced amino acid alphabet that excludes cysteine). We default to uniform probability over all possible sequence positions.

The next step in the iteration is to obtain a structure prediction corresponding to the sequence with the proposed change. This prediction provides the structural information that is used to compute the values of the individual constraint functions. These values are then combined to produce the value of the energy function, as described above. This energy function is evaluated on the overall design with and without the proposed mutation, which we denote *E*(*x*^*^) and *E*, respectively. The mutation is accepted with probability

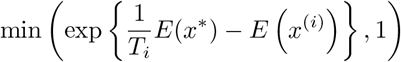

where *T*_*i*_ is the temperature parameter at iteration *i* that decays geometrically over the optimization. By default, our optimization leverages user-specified values *T*_max_ and *T*_min_, with a decay schedule given as

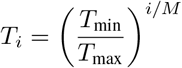

where *M* is the user-specified number of annealing steps. We report specific values for *T*_max_ and *M* in the experiment descriptions below and default to *T*_min_ = 0.0001.

#### A.1.5. Single chain constraint

Our language accommodates both single- and multi-chain design through the use of a special “single chain” constraint. By default, without this constraint applied, all terminal symbols are assumed to correspond to separate chains. When this constraint is applied to a given node, it constrains all of the terminal symbols to be part of a single chain according to the left-to-right order defined in the syntax tree. For example, consider a syntax subtree with *x*2 → *x*_3_*AB* and *x*_3_ → *CD* productions. A single chain constraint applied to node *x*_2_ would create a chain consisting of a contiguous sequence *CDAB*. Unlike other constraints, this constraint is enforced as part of structure prediction, prior to the energy function compilation.

### A.2. Constraint implementation

#### A.1.1. ESMFold structure prediction

We obtain all-atom structure predictions using ESMFold (16), where the prediction is made over the entire protein sequence and is represented as a set of atomic coordinates and their corresponding residue identities and indices. This predicted structure is the basis for the structural information passed to each of the specific constraint functions. When a constraint is defined on a subtree, that constraint only has access to the structural information (atomic coordinates, etc.) of the sequence encoded by that subtree.

#### A.2.2. Structure prediction confidence (ptm and plddt)

ESMFold produces a pTM score, which indicates the model’s confidence in the overall structure prediction, and a per-atom pLDDT score, which indicates the model’s confidence in the specific atomic coordinate prediction. The pTM value and the mean of the backbone pLDDT values are constraints that are meant to steer the optimization toward structures with higher structure prediction confidence, which is associated with naturally plausible and designable structures. We use a linear combination of the quantities 1 − pTM and 1 − pLDDT (since a higher confidence/lower energy is desirable), with user-specified weights, as the returned value of the confidence constraint.

#### A.2.3. Surface-exposed hydrophobics

The surface exposed hydrophobics constraint aims to reduce the hydrophobicity of the protein surface, where high hydrophobicity leads to protein aggregation and insolubility. We implement this constraint using the Shrake-Rupley “rolling probe” algorithm to determine the surface exposed atoms (30) as implemented in the biotite Python package version 0.35.0 (31). We then calculate the fraction of atoms involved in hydrophobic residues that are also surface exposed, and we use this fraction as the output of the constraint function.

#### A.2.4. Globularity

It is sometimes desirable to encourage a protein chain to pack into a globular structure. Our globularity constraint is implemented by computing the centroid of a set of atomic coordinates, where the globularity constraint function returns the variance of the distances from all coordinates to this centroid. Intuitively, low variance indicates that all coordinates that are largely equidistant to the centroid, which is more consistent with globular packing.

#### A.2.5. Secondary structure

The secondary structure constraint steers the energy toward user-defined secondary structure. To annotate residue secondary structure, we use the P-SEA algorithm (32) as implemented by the biotite Python package (31). This constraint function returns one minus the fraction of residues that belong to the desired secondary structure element (since a higher fraction/lower energy is desirable).

#### A.2.6. Rotational symmetry

To design symmetry, we first find it helpful to tie the sequence identities across the subsequences corresponding to the asymmetric units. The first symmetry we consider is rotational symmetry, which only consider the centroids of the immediate children of the constraint’s node; for example, rotational symmetry defined on a node *x*_1_ where *x*_1_ → *x*_2_*x*_3_*x*_4_ would only consider the centroids of the individual substructures defined by *x*_2_, *x*_3_, or *x*_4_.

Using the left-to-right order of these children in the production rule, the rotational symmetry function first computes the distances among adjacent centroids, circularly wrapping to include the distance between the first and last symbols; for example, rotational symmetry defined on *x*_1_ → *x*_2_*x*_3_*x*_4_ would compute the set of distances among pairs (*x*_2_, *x*_3_), (*x*_3_, *x*_4_), and (*x*_4_, *x*_2_). The final value returned by this constraint function is the variance among all adjacent distances; intuitively, a rotational or ring-like symmetry would have equal distances among centroids. This rotational symmetry function is adopted from that used by Wicky et al. (22).

#### A.2.7. Globular symmetry

The globular symmetry constraint is defined on the centroids of the immediate children of the constraint’s node; for example, globular symmetry defined on a node *x*_1_ where *x*_1_ → *x*_2_*x*_3_*x*_4_ would only consider the centroids of the individual substructures defined by *x*_2_, *x*_3_, or *x*_4_ (this is similar to rotational symmetry described above). The globularity symmetry function computes all pairwise distances among centroids and returns the variance of these distances. In practice, this constraint function is useful for defining symmetry that is not rotational, for example, the symmetry observed in polyhedral assemblies.

#### A.2.8. All-atom coordination

One approach to designing functional proteins is to constrain (a portion of) the protein to match the structure of a known functional site in nature. We accomplish this with an allatom coordination constraint. This constraint is first defined with respect to a list of atoms from a native protein structure (outside of our designed protein), which we denote *y*_native_. We then constrain all of the atoms in the corresponding (sub)tree to match, which we denote *y*_design_, as closely as possible, the coordination of the atoms in *y*_native_. We achieve this with two functions. The first is the constrained root mean square deviation (cRMSD),

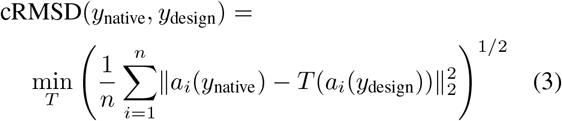

where *T* is a structural transformation, *a*_*i*_ denotes the atomic coordinates of the *i*th atom out of *n* total atoms considered, and ∥·∥ denotes a vector norm. We implement the structural alignment using the Kabsch algorithm (33) as implemented by biotite (31). The second function for constraining atomic coordination is the distance-matrix RMSD (dRMSD),

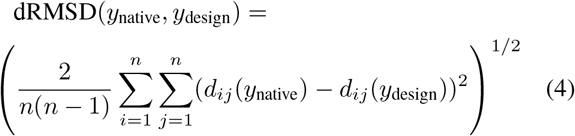

where *d*_*ij*_ is the Euclidean distance between the ith and jth atoms. The returned final value is a linear combination of the cRMSD and dRMSD values with user-specified weights. In practice, cRMSD is sometimes excluded (i.e., its weight is set to zero) in conjunction with dRMSD, as cRMSD alone does not appear sufficiently stable to create a sufficiently smooth energy landscape.

#### A.2.9. Backbone atom coordination

For a class of design tasks called fixed backbone design, we desire to only constrain the backbone atoms of the protein structure and have the optimization produce sequences that match a known backbone. This constraint is largely equivalent to the all-atom constraint described above, but rather than constraining all atoms (including side chains), this constraint is only applied to the carbon, *α*-carbon, and nitrogen atoms in the protein backbone.

#### A.2.10. Surface exposure

In some cases, we desire that a given set of residues be exposed on the surface of the protein (for example, when scaffolding a protein binding site). As with the hydrophobics constraint, we leverage the Shrake-Rupley algorithm

(30) as implemented by biotite (31). We then calculate the fraction of surface exposed atoms within the structure corresponding to the constraint’s subtree, and we use one minus this fraction as the output of the function.

#### A.2.11. Length

The length constraint requires a user-specified number of residues. In practice, we can enforce a hard length constraint by disallowing insertions and deletions during the optimization procedure, or through a function that returns increasingly high values when a sequence length goes beyond a user-specified range. In this study, whenever we apply a length constraint we take the former approach.

### A.3. Design tasks and experiments

#### A.3.1. Free hallucination

Free hallucination simply requires applying confidence and surface-exposed hydrophobic constraints to the whole protein, where we place equal weight on each term (pTM, pLDDT, and hydrophobics). In the experiments described in this study, we ran simulated annealing over 30,000 iterations with *T*_max_ = 1 across 200 seeds. We also evaluated single-sequence AlphaFold2 (18) on the final sequences produced by these 200 optimization runs.

#### A.3.2. Fixed backbone design

For fixed backbone design, we apply a weight of 2 on the dRMSD constraint, a weight of 1 on the cRMSD, pTM, and pLDDT constraints, and a weight of 0.5 on the hydrophobics constraint. As the target backbones, we used the de novo structures with PDB IDs 1QYS, 5L33, 6D0T, 6MRS, 6W3W, and 6WVS. In the experiments described in this study, we ran simulated annealing over 30,000 iterations with *T*_max_ = 1 across at least 50 seeds for each de novo backbone.

#### A.3.3. Secondary structure design

We performed protein design with partial constraints on a protein by constraining the secondary structure corresponding to different segments of the protein sequence. We place a weight of 10 on the secondary structure constraint and weights of 1 on pTM, pLDDT and hydrophobics constraints. Our programs specify the secondary structure corresponding to two discrete subsequences, where we program (1) all alpha, (2) all beta, and (3) mixed alpha and beta secondary structure. We ran simulated annealing over 30,000 iterations *T*_max_ = 1 for 10 seeds for each of these three programs (30 optimization trajectories in total).

#### A.3.4. Single functional site scaffolding

To program functional site scaffolding on a de novo backbone, we divide a single-chain sequence into three segments: a sequence in the middle segment (with an all-atom coordination constraint and a surface exposure constraint) flanked by two “free” sequences. pTM, pLDDT, and hydrophobics constraints are also applied to the full protein. We apply a weight of 2 to the cRMSD and dRMSD constraints, and a weight of 1 to the pTM, pLDDT, and hydrophobics constraints.

We attempted to scaffold five protein binding sites, the first three of which were successfully scaffolded by Wang et al. (20):

1. IL10: We used the residue indices 31–40, inclusive, of chain L in the PDB structure 1Y6K, corresponding to the IL10 binding site of IL-10R1 (34).
2. ACE2: We used the residue indices 5–23, inclusive, of chain A in the PDB structure 6M0J, corresponding to the ACE2 binding site of the SARS-CoV-2 spike receptor binding domain (RBD) (35).
3. C3d: We used the residue indices 104–126 and 170– 184, inclusive, of chain A in the PDB structure 1GHQ, corresponding to the C3d binding site of complement receptor 2 (36).
4. HA2: We used the residue indices 14–21, 33–42, and 45–49, inclusive, of chain B in the PDB structure 5JW3, corresponding to the influenza HA2 epitope of the antibody MEDI8825 (37).
5. RBD: We used the residue indices 439–450 and 498– 506, inclusive, of chain C in the PDB structure 7MMO, corresponding to the SARS-CoV-2 RBD epitope of the antibody bebtelovimab (38).

We ran simulated annealing over 30,000 iterations with *T*_max_ = 1 for 1,000 seeds for each of the five binding sites (5,000 optimization trajectories in total).

#### A.3.5. Symmetric and homo-oligomer design

We first program single-chain symmetry using a rotational symmetry constraint applied to the top-level node. In our program, we also tie the sequences across the subsequences corresponding to the asymmetric units such that we only use a single terminal symbol; an example program for designing 3-fold symmetry is provided in Figure 3A. We place a weight of 1 on the symmetry constraint, as well as weights of 1 on pTM, pLDDT, and hydrophobics constraints. We also place length constraints on the terminal nodes. We specify programs where we increase the fold-symmetry from 3-to 8-fold. We also vary the lengths that constrain the terminal symbol such that the full sequence has approximately 200, 300, or 400 residues (for example, a 200-residue protein with 3-fold symmetry would have length constraints of 66 on its terminal symbols). We ran simulated annealing over 30,000 iterations with a starting temperature of 1 for 10 seeds for each of the six fold symmetries and each of the three length constraints (for a total of 180 optimization trajectories).

We also designed larger homo-oligomers similarly, but removing the single-chain constraint from the top-level node. We designed trimeric, tetrameric, hexameric, and octameric homo-oligomers with a globular symmetry constraint applied to the top level node. We placed a weight of 1 on the symmetry constraint, as well as weights of 1 on pTM, pLDDT, and hydrophobics constraints. We also placed a weight of 0.1 on globularity constraints that are applied to each terminal symbol. We applied length constraints such that the full complex contained 720 residues (for example, the hexamer would consist of length-120 protomers). We ran simulated annealing over 30,000 iterations with *T*_max_ = 1 for 10 seeds for each of oligomerization levels (for a total of 40 optimization trajectories).

#### A.3.6. Two-level symmetry design

We program two levels of symmetry using the productions

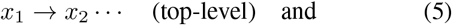

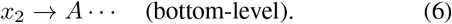

In these programs, we place the single-chain constraint on *x*_2_, so the final designs are protein homo-oligomers. We place a globularity symmetry constraint on *x*_1_; to control the top-level symmetry, we repeat *x*_2_ according to the desired oligomerization. We place a rotational symmetry constraint on *x*_2_; to control the bottom-level symmetry, we repeat *A* according to desired fold symmetry. We also place pTM, pLDDT, and hydrophobics constraints on the full protein; we place globularity constraints on *x*_2_. We compile constraints into an energy function with weights of 1 on all terms.

We enumerated programs over the grid varying both the top and bottom levels of symmetry from 2 to 4. We constrained lengths to 200 residues in total for the dimer of 2-fold; length-250 for the dimer of 3-fold; length-400 for the dimer of 4-fold, the trimer of 2-fold, the trimer of 3-fold, and the tetramer of 2-fold; length-450 for the trimer of 4-fold and the tetramer of 3-fold; and length-500 for the tetramer of 4-fold. We ran simulated annealing over 30,000 iterations with *T*_max_ = 1 for 10 seeds for each of these programs (for a total of 90 optimization trajectories).

#### A.3.7. Structural novelty

We quantify a given design for structural novelty by running an exhaustive search over the PDB version 2022-08 (http://www.rcsb.org/) (39) to find the experimental structure with the highest TM-score to the designed structure, normalizing by the designed structure length, using TM-align version 20210107 (40).

#### A.3.8. Inverse folding roundtrip experiments

We assessed the “designability” of a structure prediction produced by our optimization procedure by “roundtripping” the protein through an inverse folding model. More specifically, given a predicted structure from our optimization loops, we first use ESM-IF1 (21), an independently trained inverse folding model, to sample 10 sequences with temperature 0.1 from the backbone coordinates. We then run these sequences through ESMFold and compute the cRMSD between the starting and the roundtripped backbone atoms of the predicted structure.

We performed this roundtrip experiment for 1,000 predicted structures that were obtained by first uniformly sampling one of the 180 symmetric single-chain optimization trajectories and then uniformly sampling one of the intermediate structure predictions within a given design loop (i.e., we do not restrict this analysis to the best pLDDT structure over a design loop, which are highly biased toward high pLDDTs). We report the relationship between ESM-IF1 perplexities of all sample structures and the corresponding cRMSD values. We also report the relationship between the pLDDT of the starting structure and the minimum RMSD over the structure for the 10 inverse-folded sequences. We also repeated the same experiment for 1,000 predicted structures that were obtained by first uniformly sampling over the 90 two-level symmetry optimization trajectories and then uniformly sampling one of the intermediate structure predictions within a given design loop. We also report the same metrics as in the single-chain evaluation.

#### A.3.9. Symmetric functional site scaffolding

We designed proteins that symmetrically scaffold multiple functional sites by using the tree described for the singlesite functional scaffold but replicating it according to the desired fold symmetry and adding a rotational symmetry constraint to the top-level node; an example program for a 3-fold functional site scaffold can be found in Figure 5A. We use weights of 10 on the cRMSD and dRMSD constraints and weights of 1 on the pTM, pLDDT, rotational symmetry, binding site surface exposure, and hydrophobics constraints. We ran simulated annealing over 30,000 iterations with a starting temperature of 1 over 20 seeds for the design of 3-fold scaffolds of the IL10 and ACE2 binding sites described above, as well as 20 seeds for the design of 5-fold scaffolds of the ACE2 binding site (for a total of 60 optimization loops).

#### A.3.10. Hierarchical asymmetric symmetry design

We increased the level of hierarchical complexity in our programs by designing with three levels of constraints. The top level specifies two asymmetric subunits. Each asymmetric subunit itself has two-level symmetry (similar to the setting described above): we specifically consider the dimer of 2-fold (2×2), the dimer of 3-fold (2×3), and the trimer of 2-fold (3×2). We write programs consisting of (1) two asymmetric 2×2s complexed together, (2) a 2×2 and a 3×2 complexed together, and (3) a 2×2 and a 2×3 completed together. An example program for the two asymmetric 2×2s is provided in Figure 5C. We use weights of 1 on all constraints (pTM, pLDDT, hydrophobics, rotational/globular symmetry, and globularity). We ran simulated annealing over 30,000 iterations with *T*_max_ = 1 over 10 seeds for each of the three programs described above (for a total of 30 optimization loops).

**Figure S1.**
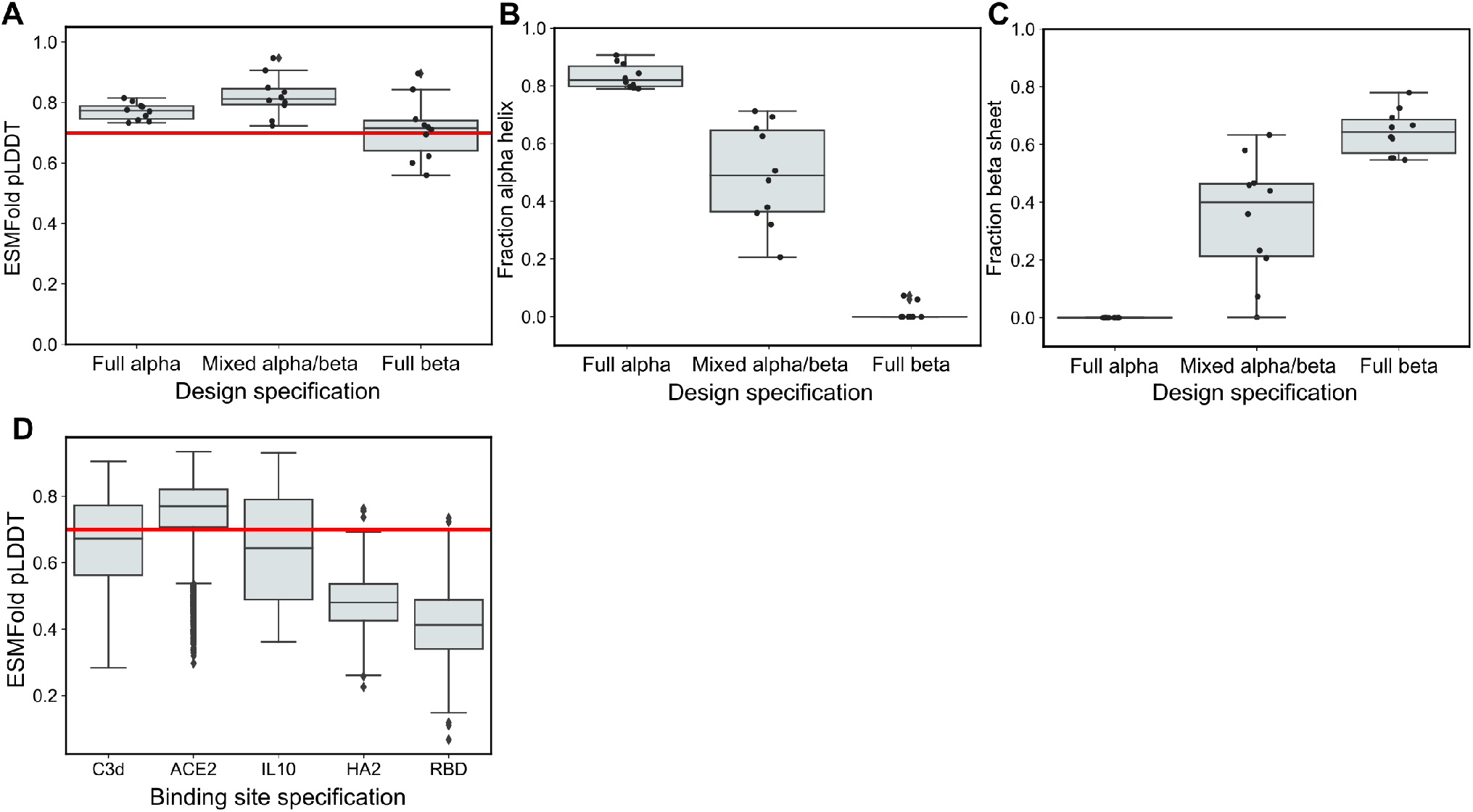
Additional plots for secondary structure design and functional site scaffolding. **(A)** ESMFold pLDDT values for different secondary structure design specifications (10 seeds per specification). A red line is plotted at pLDDT = 0.7. **(B)** The fraction of residues that are part of alpha helices for different secondary structure design specifications (10 seeds per specification). **(C)** The fraction of residues that are part of beta sheets for different secondary structure design specifications (10 seeds per specification). **(D)** ESMFold pLDDT values for different functional site scaffolding design runs (1,000 seeds per binding site).

**Figure S2.**
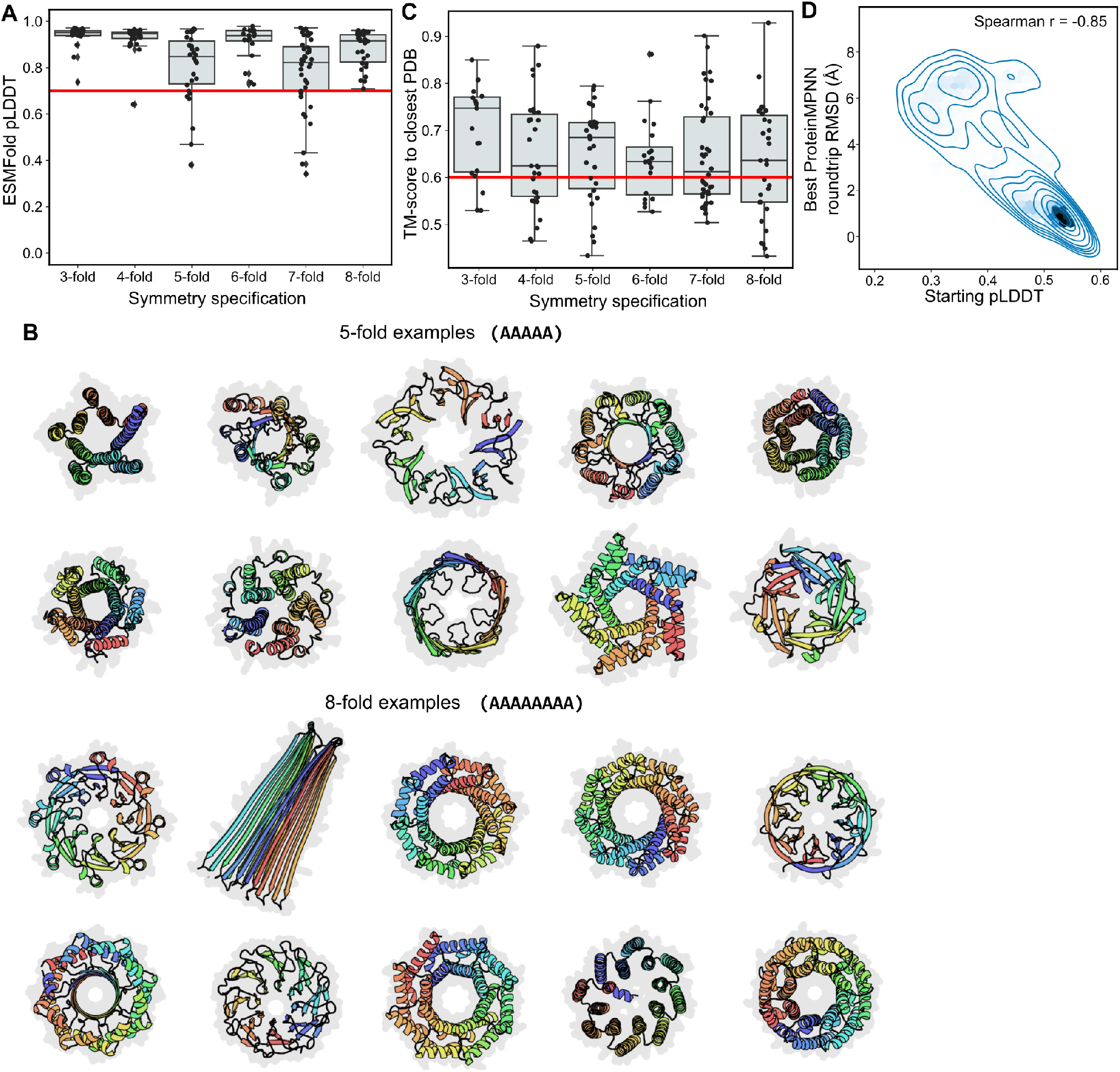
Additional plots for the design of symmetric single chains. **(A)** ESMFold pLDDT values for different fold symmetry design specifications (30 seeds per specification). A red line is plotted at pLDDT = 0.7. **(B)** Ten randomly sampled designs for the design of 5- and 8-fold symmetry. **(C)** TM-scores for different fold symmetry design specifications (30 seeds per specification); the TM-score is between the best design and the closest structure in the PDB. A red line is plotted at TM-score = 0.6. **(D)** Samples obtained by inverse folding with ProteinMPNN. On the *x*-axis is the pLDDT of the designed structure prior to the roundtrip and on the *y*-axis is the roundtrip RMSD.

**Figure S3.**
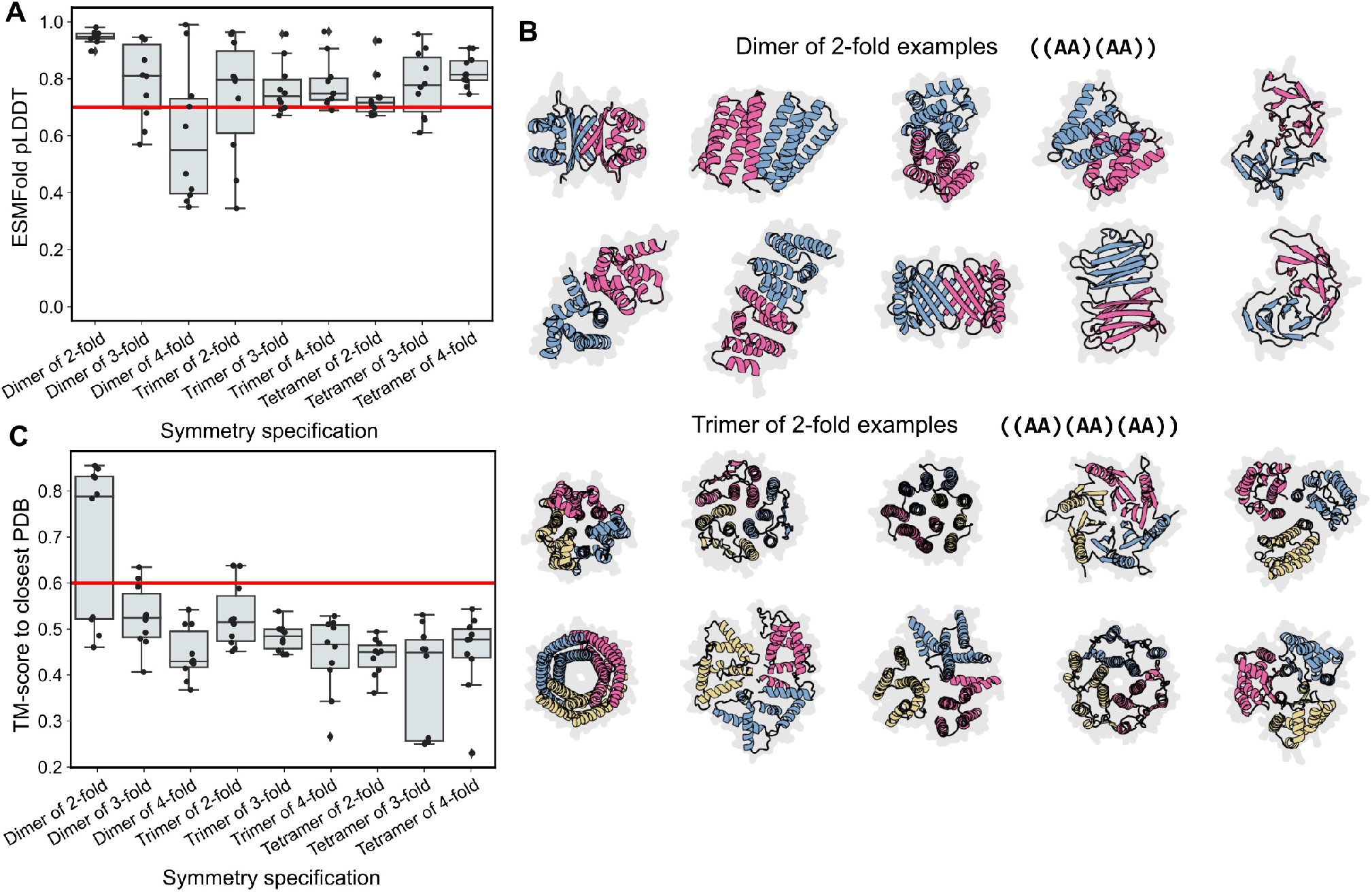
Additional plots for the design of two-level symmetry. **(A)** ESMFold pLDDT values for different two-level symmetry design specifications (10 seeds per specification). A red line is plotted at pLDDT = 0.7. **(B)** All ten of the designs of a dimer of 2-fold and of a trimer of 2-fold symmetry. **(C)** TM-scores for different two-level symmetry design specifications (10 seeds per specification); the TM-score is between the best design and the closest structure in the PDB. A red line is plotted at TM-score = 0.6.

**Figure S4.**
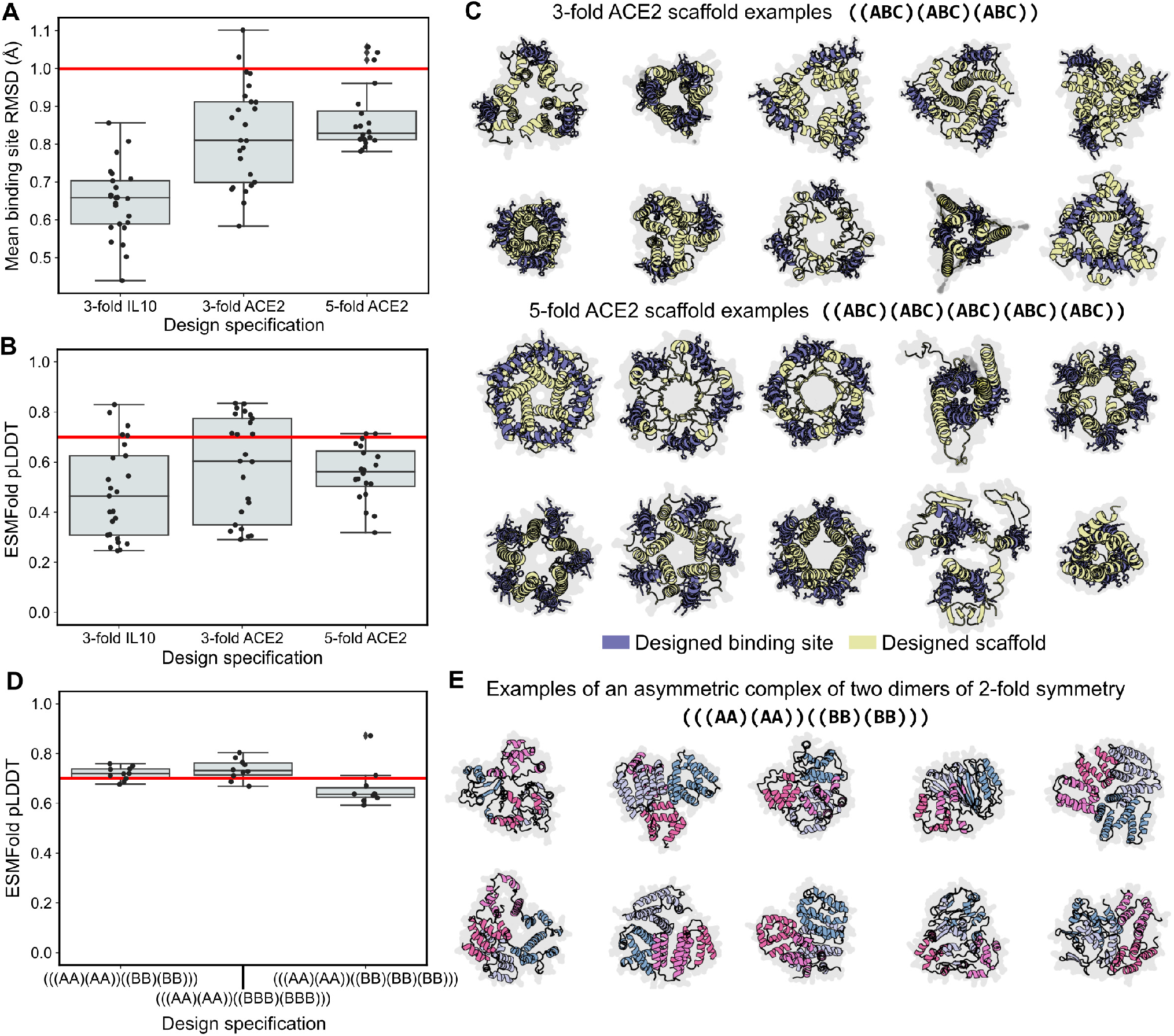
Additional plots for multi-level hierarchical design. **(A)** ESMFold pLDDT values for different binding site scaffolds with fold symmetry (20 seeds per specification). A red line is plotted at pLDDT = 0.7. **(B)** RMSD values for different binding site scaffolds with fold symmetry (20 seeds per specification). The mean RMSD is reported across either three or five binding sites. A red line is plotted at pLDDT = 0.7. **(C)** Ten randomly sampled designs for the design of 3- and 5-fold symmetric ACE2 binding site scaffolds. **(D)** ESMFold pLDDT values for different asymmetric-symmetric design specifications (10 seeds per specification). A red line is plotted at pLDDT = 0.7. (E) All ten of the designs of an asymmetric complex of two dimers of 2-fold symmetry.

